# Novel orthonairovirus in rodents and shrews, Gabon

**DOI:** 10.1101/2022.01.22.477371

**Authors:** Takehiro Ozeki, Haruka Abe, Yuri Ushijima, Chiméne Nze-Nkogue, Etienne F Akomo-Okoue, Ghislain W.E Ella, Lilian B.M Koumba, Branly C.B.B Nzo, Rodrigue Mintsa-Nguema, Patrice Makouloutou-Nzassi, Boris K Makanga, Fred L.M Nguelet, Georgelin N Ondo, Marien J.V.M Mbadinga, Yui Igasaki, Sayaka Okada, Bertrand Lell, Laura C. Bonney, Roger Hewson, Yohei Kurosaki, Jiro Yasuda

## Abstract

Small mammals harbor various zoonotic viruses and are natural reservoirs for emerging viruses. Here, we identified a novel orthonairovirus, which is genetically close to the virus suggested the association with human neural diseases. The virus was found in 24.6% of the small mammals captured in Gabon, Central Africa.

## Text

Small mammals, including rodents and shrews, are natural reservoirs of many zoonotic pathogens. These animals host highly pathogenic human viruses such as Lassa, Machupo, Junin, hanta, and Crimean-Congo hemorrhagic fever viruses, all of which belong to order *Bunyavirales* (*1*). Rodent-borne viral diseases, such as Lassa fever, have emerged in sub-Saharan Africa. Thus, local residents are at a potentially high risk of viral exposure under poor sanitary conditions, particularly in rural areas (*2*). Moreover, several viruses, including Tanganya virus of genus *Orthohantavirus* and Thiafora virus (TFAV) of genus *Orthonairovirus*, both belonging to *Bunyavirales*, have been identified in the musk shrews (*Crocidura* sp.) in Western Africa (*3,4*). Accordingly, small mammal populations hosting numerous zoonotic viruses may pose a public health threat to humans in sub-Saharan Africa.

Due to high rate of annually reported Lassa fever cases in Western Africa, several investigations of small mammal-borne viruses have been conducted in the region, thereby multiple zoonotic viruses have been identified (*5–8*). In contrast, in Central Africa, no clinical cases of Lassa fever or other rodent-borne human viral diseases have been reported which indicates that few small mammal-borne viral surveillance studies have been conducted (*5*). Nonetheless, recent serological surveillance studies have provided evidence of past infection of rodent-borne viruses among residents of Central African countries (*9,10*). This suggested that unrecognized zoonotic viruses might be present in this region. Therefore, to understand the potential risks of transmission of known and unknown zoonotic small mammal borne viruses, we investigated the viruses present among small mammal populations in Gabon, Central Africa.

### The Study

Between 2019 and 2020, 281 animals (152 rodents and 129 shrews) were captured using Sherman and Tomahawk traps placed in a forest near the suburban area, and bushes around human dwellings in Lambaréné, Central Gabon. After organ specimens were collected by dissection, tissue RNA was extracted from kidney homogenates, as described previously (*10*). Virus screening was performed using the PrimeScript II High Fidelity One Step RT-PCR Kit (Takara Bio, Shiga, Japan). We initially targeted partially conserved nucleotide sequences of three genera of viruses: *Mammarenavirus, Orthohantavirus*, and *Orthonairovirus*, belonging to the order *Bunyavirales*. For the RT-PCR detection of mammarenavirus and orthohantavirus, we employed the pan-viral family primer sets described previously (*8,11*). For orthonairovirus, a primer set was designed based on an alignment of sequences of a highly conserved region on the large segment (Appendix Table). All RT-PCR was performed under the following conditions: 10 min at 45 °C, 2 min at 94 °C, 35 cycles each of 10 s at 98 °C, 15 s at 45 °C, and 10 s at 68 °C. All amplicons were confirmed by Sanger sequencing, and the obtained sequences were identified by BLAST search (https://blast.ncbi.nlm.nih.gov).

The species of animals captured in this study were identified by analyzing their cytochrome b gene nucleotide sequences, as described previously (*10*). The animal species were determined by BLAST search after performing Sanger sequencing.

After initial screening, viral sequences of mammarenavirus and orthohantavirus were not detected in any of the samples (Table 1). By contrast, novel orthonairovirus-like sequences were detected in 69 out of the 281 sampled animals (24.6%). BLAST search revealed that the newly identified sequences showed high similarity to Erve virus (ERVEV) or TFAV (62.8-74.4%). These viruses were previously identified among the *Crocidura* sp. found in France and Senegal, respectively (*4,12*). Notably, the virus prevalence in *Crocidura* sp. was significantly higher than that in all other captured rodents (virus prevalence in *Crocidura* sp. vs. rodents: 34.1% (44/129), odds ratio: 2.63, 95% CI: 1.50-4.62, *p* <0.001). This novel orthonairovirus was named Lamusara virus (LMSV), based on the identified place (Lambaréné) and virus hosts (“musaraigne”; “shrew” in French and “ra”; “rodent” in French).

**Table 1.** Species of captured small mammals and results of virus screening

| Mammal species | Common name | *Mammarenavirus* | *Orthohantavirus* | *Orthonairovirus* |
| --- | --- | --- | --- | --- |
| *Crocidura goliath* | Goliath shrew | 0/128* | 0/128 | **44/128** |
| *Crocidura poensis* | Fraser's musk shrew | 0/1 | 0/1 | 0/1 |
| *Hybomys univittatus* | Peter's striped mouse | 0/3 | 0/3 | 0/3 |
| *Hylomyscus* sp. | African wood mouse | 0/2 | 0/2 | 0/2 |
| *Lemniscomys striatus* | Typical striped grass mouse | 0/3 | 0/3 | **1/3** |
| *Lophuromys* sp. | Brush-furred mouse | 0/7 | 0/7 | 0/7 |
| *Mus minutoides* | African pygmy mouse | 0/41 | 0/41 | **6/41** |
| *Mus musculus* | House mouse | 0/3 | 0/3 | 0/3 |
| *Oenomys hypoxanthus* | Common rufous-nosed rat | 0/6 | 0/6 | **2/6** |
| *Praomys misonnei* | Misonne's soft-furred mouse | 0/65 | 0/65 | **11/65** |
| *Rattus rattus* | Black rat | 0/16 | 0/16 | **5/16** |
| *Stochomys longicaudatus* | Target rat | 0/6 | 0/6 | 0/6 |
|  |  | 0/281 | 0/281 | 69/281 |

\*Number of virus-positive individuals/number of captured animals. Positive number is indicated in boldface.

Genus *Orthonairovirus* belongs to family *Nairoviridae*, which includes enveloped and negative-sense single-stranded RNA viruses (*13*). Its genome comprises large, medium, and small segments encoding the RNA-dependent RNA polymerase (L), glycoprotein precursor (GPC), and nucleoprotein (N), respectively (*14*).

To determine longer viral genome sequences, we performed RT-PCR using RNA samples positive for the LMSV genome. To design the deduced primer sets, the nucleotide sequences of each genome segment of ERVEV and TFAV were aligned (GenBank accession numbers: JF911697-JF911699, KU925458-KU925460 (ERVEV), and NC_039220-NC_039222 (TFAV); Appendix Table). Once the amplicons were confirmed as the LMSV genome, Sanger sequencing was repeatedly performed to close sequence gaps.

Whole genome sequences of LMSV, consisting of three segments: large, medium, and small, were successfully determined (DDBJ accession numbers: LC671712-LC671787) in accordance with orthonairovirus genome organization. Each genome segment comprises a single open reading frame (ORF) encoding the L (11,583 nt / 3,861 aa), GPC (3,819 nt / 1,273 aa), and N (2,016 nt / 672 aa) proteins, respectively. Phylogenetic analysis was also performed among orthonairovirus protein-coding region of nucleotide sequences using IQ-TREE (http://iqtree.cibiv.univie.ac.at/) to genetically characterize LMSV. The results showed that all three LMSV encoding proteins were phylogenetically close to ERVEV and TFAV, yet formed unique phylogenetic clusters that differed from other orthonairovirus (Figure). Moreover, there were at least two distinct genotypes in the LMSV cluster of each segment according to the constructed trees. Comparison of nucleotide and amino acid sequences between two LMSV genotypes (strains CG002 vs. CG020) revealed that the sequences of the GPC protein were highly conserved (82.5% and 92.6% at nucleotide and amino acid levels, respectively), whereas the sequences of L and N proteins were relatively divergent between the two genotypes (L: 68.6% and 66.9% at nucleotide and amino acid levels, respectively; N: 59.9% and 61.4% at nucleotide and amino acid levels, respectively) (Table 2).

**Figure.**
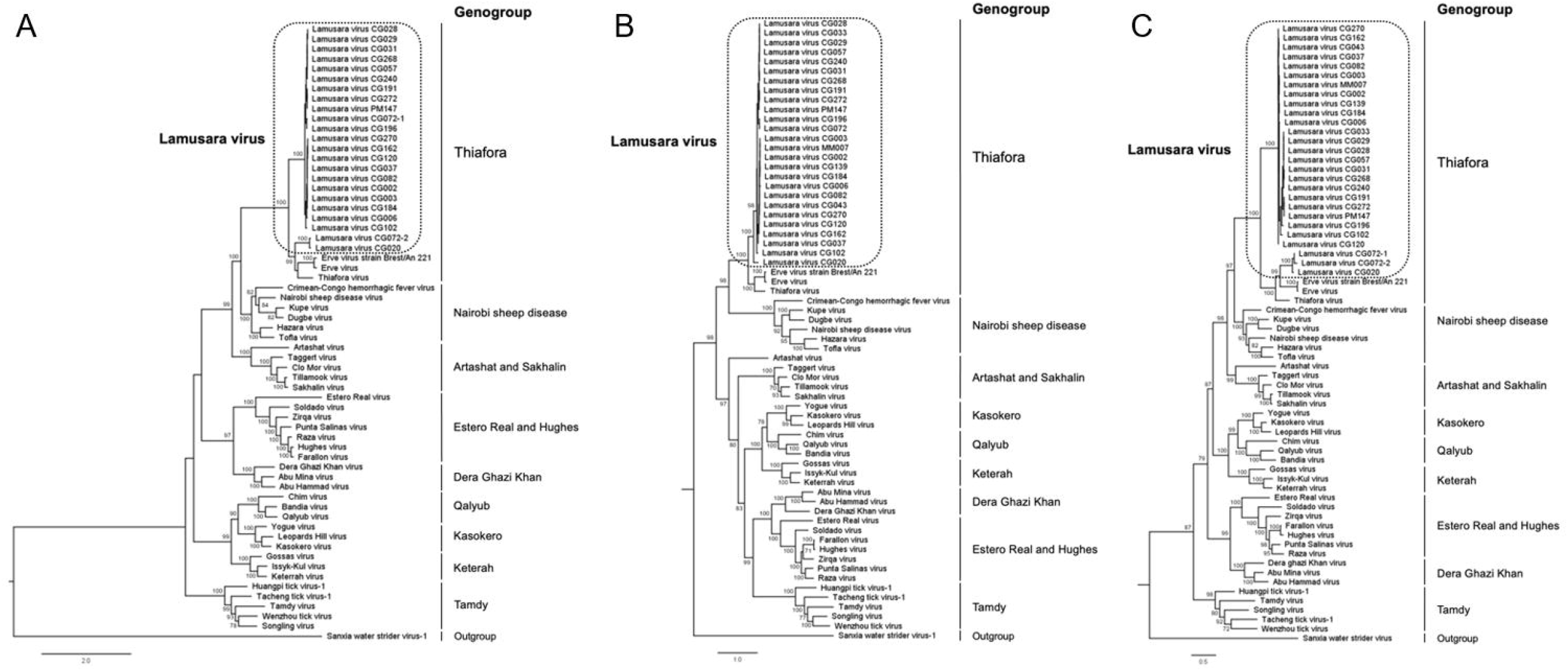
Maximum-likelihood phylogenetic trees of LMSV were constructed based on L protein partial nucleotide sequences (at position 6,913-11,471 nt on LMSV large-segment (DDBJ accession number: LC671712)) (A), GPC complete nucleotide sequences (B) and N protein complete nucleotide sequences (C) using IQ-tree (http://iqtree.cibiv.univie.ac.at/) with 1,000 bootstraps. Bootstrap values of ≧70% are shown at the nodes of the trees. Nine genogroups of genus *Orthonairovirus* are shown on the right of the trees (*4*) and LMSV clusters in each segment were circled by broken line. The scale bar indicates nucleotide substitutions per site.

**Table 2.**
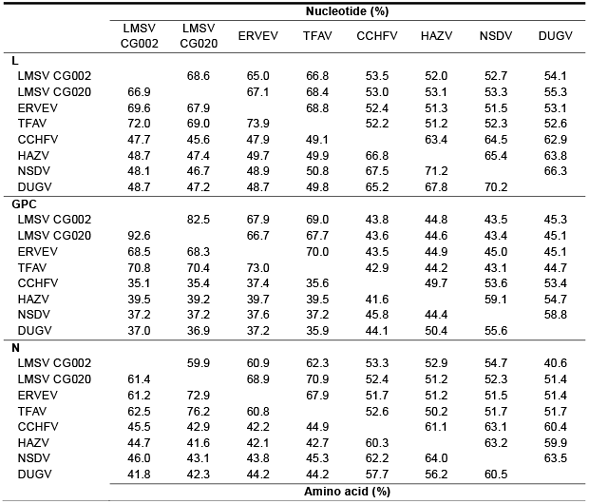
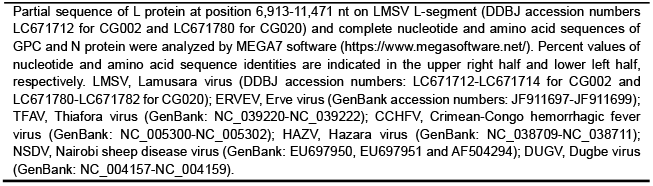
Pairwise comparisons of nucleotide and amino acid sequences among representative orthonairoviruses

|  | Nucleotide (%) |  |  |  |  |  |  |  |
| --- | --- | --- | --- | --- | --- | --- | --- | --- |
|  | LMSV<br>CG002 | LMSV<br>CG020 | ERVEV | TFAV | CCHFV | HAZV | NSDV | DUGV |
| **L** |  |  |  |  |  |  |  |  |
| LMSV CG002 |  | 68.6 | 65.0 | 66.8 | 53.5 | 52.0 | 52.7 | 54.1 |
| LMSV CG020 | 66.9 |  | 67.1 | 68.4 | 53.0 | 53.1 | 53.3 | 55.3 |
| ERVEV | 69.6 | 67.9 |  | 68.8 | 52.4 | 51.3 | 51.5 | 53.1 |
| TFAV | 72.0 | 69.0 | 73.9 |  | 52.2 | 51.2 | 52.3 | 52.6 |
| CCHFV | 47.7 | 45.6 | 47.9 | 49.1 |  | 63.4 | 64.5 | 62.9 |
| HAZV | 48.7 | 47.4 | 49.7 | 49.9 | 66.8 |  | 65.4 | 63.8 |
| NSDV | 48.1 | 46.7 | 48.9 | 50.8 | 67.5 | 71.2 |  | 66.3 |
| DUGV | 48.7 | 47.2 | 48.7 | 49.8 | 65.2 | 67.8 | 70.2 |  |
| **GPC** |  |  |  |  |  |  |  |  |
| LMSV CG002 |  | 82.5 | 67.9 | 69.0 | 43.8 | 44.8 | 43.5 | 45.3 |
| LMSV CG020 | 92.6 |  | 66.7 | 67.7 | 43.6 | 44.6 | 43.4 | 45.1 |
| ERVEV | 68.5 | 68.3 |  | 70.0 | 43.5 | 44.9 | 45.0 | 45.1 |
| TFAV | 70.8 | 70.4 | 73.0 |  | 42.9 | 44.2 | 43.1 | 44.7 |
| CCHFV | 35.1 | 35.4 | 37.4 | 35.6 |  | 49.7 | 53.6 | 53.4 |
| HAZV | 39.5 | 39.2 | 39.7 | 39.5 | 41.6 |  | 59.1 | 54.7 |
| NSDV | 37.2 | 37.2 | 37.6 | 37.2 | 45.8 | 44.4 |  | 58.8 |
| DUGV | 37.0 | 36.9 | 37.2 | 35.9 | 44.1 | 50.4 | 55.6 |  |
| **N** |  |  |  |  |  |  |  |  |
| LMSV CG002 |  | 59.9 | 60.9 | 62.3 | 53.3 | 52.9 | 54.7 | 40.6 |
| LMSV CG020 | 61.4 |  | 68.9 | 70.9 | 52.4 | 51.2 | 52.3 | 51.4 |
| ERVEV | 61.2 | 72.9 |  | 67.9 | 51.7 | 51.2 | 51.5 | 51.4 |
| TFAV | 62.5 | 76.2 | 60.8 |  | 52.6 | 50.2 | 51.7 | 51.7 |
| CCHFV | 45.5 | 42.9 | 42.2 | 44.9 |  | 61.1 | 63.1 | 60.4 |
| HAZV | 44.7 | 41.6 | 42.1 | 42.7 | 60.3 |  | 63.2 | 59.9 |
| NSDV | 46.0 | 43.1 | 43.8 | 45.3 | 62.2 | 64.0 |  | 63.5 |
| DUGV | 41.8 | 42.3 | 44.2 | 44.2 | 57.7 | 56.2 | 60.5 |  |
|  |  |  |  | Amino acid (%) |  |  |  |  |

Partial sequence of L protein at position 6,913-11,471 nt on LMSV L-segment (DDBJ accession numbers LC671712 for CG002 and LC671780 for CG020) and complete nucleotide and amino acid sequences of GPC and N protein were analyzed by MEGA7 software (https://www.megasoftware.net/). Percent values of nucleotide and amino acid sequence identities are indicated in the upper right half and lower left half, respectively. LMSV, Lamusara virus (DDBJ accession numbers: LC671712-LC671714 for CG002 and LC671780-LC671782 for CG020); ERVEV, Erve virus (GenBank accession numbers: JF911697-JF911699); TFAV, Thiafora virus (GenBank: NC\_039220-NC\_039222); CCHFV, Crimean-Congo hemorrhagic fever virus (GenBank: NC\_005300-NC\_005302); HAZV, Hazara virus (GenBank: NC\_038709-NC\_038711); NSDV, Nairobi sheep disease virus (GenBank: EU697950, EU697951 and AF504294); DUGV, Dugbe virus (GenBank: NC\_004157-NC\_004159).

LMSV showed the highest sequence similarity to TFAV compared to other orthonairovirus, at nucleotide and amino acid levels in all three ORFs (LMSV CG002 vs. TFAV; L: 66.8% and 72.0% at nucleotide and amino acid levels, respectively; GPC: 69.0% and 70.8% at nucleotide and amino acid levels, respectively; N: 62.3% and 62.5% at nucleotide and amino acid levels, respectively) (Table 2). According to the criteria described by Walker et al., LMSV should be grouped into the genogroup *Thiafora* based on the amino acid sequence homology value (>52%) of the N protein between LMSV, ERVEV, and TFAV (*4*).

## Conclusions

In this study, we identified a novel orthonairovirus, LMSV, drawn from small mammals captured in Gabon, Central Africa. According to our results, LMSV should be assigned to the genogroup *Thiafora*, based on its genetic relationship with ERVEV and TFAV. Further, we found that *Crocidura* sp. could be considered as the main host for LMSV. We also demonstrated that LMSVs have acquired significant sequence diversity. These results suggest the unique evolution and host adaptation of LMSV circulating among wildlife in Gabon.

A previous study suggested that ERVEV has a potential risk of causing neuropathogenic human diseases including thunderclap headaches (*15*). Here, we demonstrated preliminary evidence for the presence of possible causative agents of zoonotic diseases in Gabon. However, further studies are needed to understand the biological characteristics of LMSV. Moreover, surveillance studies of the seroprevalence of rodent-borne viruses, such as mammarenavirus and orthohantavirus, have previously been conducted among local Gabonese residents (*9,10*). By contrast, there are no reports specifically targeting seroprevalence of orthonairovirus. Taken together, our findings would provide novel insight into small mammal borne virus related to public health issues in Gabon.

## Acknowledgements

We thank Ms. Miku Takano (Nagasaki University) and Ms. Izumi Suzumori (Japan International Cooperation Agency (JICA)) for the management of logistics, material transportation, and linguistic support. We are also grateful to all staff in the Department of Emerging Infectious Diseases of Nagasaki University, CERMEL, IRET, and JICA for their support and considerable encouragement. We would like to thank Editage (www.editage.com) for English language editing. This work was supported by the Science and Technology Research Partnership for Sustainable Development (SATREPS) from the Japan International Cooperation Agency (JICA) and the Japan Agency for Medical Research and Development (AMED), JP21jm0110013, AMED, JP21jm0210072, and the Japan Society for the Promotion of Science (KAKENHI), JP17KK0170, and JP21K10415.

## Author’s Bio

Mr. Takehiro Ozeki is a graduate student at Nagasaki University. His research interests include emerging infectious diseases, virus discovery, and host innate immune defense.

---

**Appendix Table.**
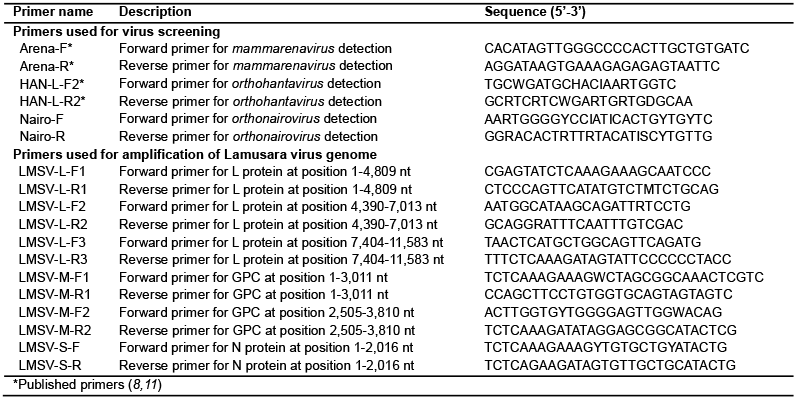
Primers used for virus screening and amplification of Lamusara virus genome

## Notes

### Competing Interest Statement

The authors have declared no competing interest.

